# GenDiS3 database: census on prevalence of protein domain superfamilies of known structure in the entire sequence database

**DOI:** 10.1101/2025.03.14.643262

**Authors:** Sarthak Joshi, Shailendu Mohapatra, Dhwani Kumar, Adwait Joshi, Meenakshi Iyer, Ramanathan Sowdhamini

## Abstract

Despite the vast amount of sequence data available, a significant disparity exists between the number of protein sequences identified and the relatively few structures that have been resolved. This disparity highlights the challenge in structural biology to bridge the gap between sequence information and three-dimensional structural data, and the necessity for robust databases capable of linking distant homologues to known structures. Studies have indicated that there is a limited number of structural folds, despite the vast diversity of proteins. Hence, computational tools can enhance our ability to classify protein sequences, much before their structures are determined or their functions characterised, thereby bridging the gap between sequence and structural data. GenDiS is a repository with information on Genomic Distribution of protein domain superfamilies, involving a one-time computational exercise to search for trusted homologues of protein domains of known structures against the vast sequence database. We have updated this database employing advanced bioinformatics tools, including DELTA-BLAST for initial detection of hits and HMMSCAN for validation, significantly improving the accuracy of domain identification. Using these tools, over 151 million sequence homologues for 2060 superfamilies (SCOPe) were identified and 116 million out of them were validated as true positives. Through a case study on glycolysis-related enzymes, variations in domain architectures of these enzymes are explored, revealing evolutionary changes and functional diversity amongst these essential proteins. We present another case, LoG gene, where there one can tune-in and find significant mutations across the evolutionary lineage. The GenDiS database, GenDiS3, and the associated tools made available at https://caps.ncbs.res.in/gendis3/, offer a powerful resource for researchers in functional annotation and evolutionary studies.

Database URL: https://caps.ncbs.res.in/gendis3/

## Introduction

Proteins, the fundamental building blocks of biological functions, are composed of one or more specific functional regions known as domains. Each domain within a protein can be characterised by its distinct structural and functional properties. Structurally, domains are classified by databases such as SCOP (Structural Classification of Proteins) (1,2) and CATH (Class, Architecture, Topology, and Homologous superfamily) (3,4), which organise protein structures based on hierarchical relationships of folding patterns and architecture. On the sequence level, domains are defined through databases like Pfam (5) and InterPro (6), which use amino acid sequences to identify and group protein domains across different species.

Despite the extensive sequence data that has been accumulated, there remains a significant disparity between the sheer number of protein sequences identified and the relatively few protein structures that have been resolved. This gap underscores a major challenge in structural biology: the need to connect sequence information with three-dimensional structural understanding. Addressing this challenge is critical, as it enhances our comprehension of protein function and interactions. Therefore, there is a pressing need for robust and comprehensive databases that can accurately link distant homologues to known structures, facilitating the prediction and modelling of protein structures from sequence data alone.

Research has shown that there is a limited number of structural folds, despite the immense diversity of protein sequences (7,8). Although the Protein Data Bank (PDB) (9) has experienced substantial growth in the number of deposited structures, the discovery of new folds has not kept pace with this increase. The relatively stable number of unique folds suggests that protein structural diversity is constrained compared to the exponential growth of sequence data. This observation underscores the importance of computational methods like Hidden Markov Models (HMMs) and Position-Specific Scoring Matrices (PSSMs) in protein classification. These methods can leverage structural conservation to enhance detection sensitivity, especially when classifying protein domains with low sequence identity among homologous proteins. Basic tools like BLAST (10) often fall short in identifying remote homologues due to subtle sequence similarities. However, advanced methods such as HMMs and PSSMs have demonstrated greater efficacy. HMMs, in particular, are adept at modelling the probabilistic nature of amino acid substitutions and insertions/deletions, providing a powerful means to detect distant relationships within protein families (11).

The previous versions of the GenDiS (Genomic Distribution of Superfamilies) database have identified homologues of SCOP superfamily members, leveraging the PASS2 database (12), which provides curated alignments for SCOP superfamily members. PASS2 is a database that provides structure-based sequence alignments for SCOPe domains that have less than 40% sequence identity with each other. PASS2 also provides Hidden Markov models derived from these alignments. The first version of GenDiS (13) utilised PSI-BLAST (14) and HMMSEARCH (15) for homologue identification. The second version (16,17) utilised CS_BLAST (18) along with PSI-BLAST. CS-BLAST enhances the sensitivity of homologue detection by taking into account the context of amino acid substitutions, thereby providing better detection of distant relationships (19), but is inherently quite greedy. The combination of CS-BLAST and PSI-BLAST, accompanied by strict validation, allowed for a more robust identification process, enabling the detection of homologues with higher sensitivity and specificity (16). Both versions utilised members from the PASS2 database, which provides high-quality alignments of SCOP superfamily members. This database is crucial for ensuring the accuracy and relevance of the homologue identification process during validation, as it offers a reliable reference for structural and functional annotation of protein sequences. The current version of the GenDiS database utilises DELTA-BLAST (Domain Enhanced Lookup Time Accelerated BLAST) (20), which integrates domain information from conserved domain databases into the BLAST search process. This allows for more sensitive and accurate detection of distant homologues by leveraging domain-specific information. For validation, the new version employs HMMSCAN, a tool from the HMMER suite (hmmer.org). This combination of DELTA-BLAST for initial detection and HMMSCAN for validation ensures a high level of accuracy and reliability in the identification process. This updated approach builds on the strengths of previous versions by incorporating more sophisticated search and validation tools, thereby improving the comprehensiveness and precision of the GenDiS database.

## Methods

### Datasets and tools used in sequence search

#### Datasets

SCOPe (Structural Classification of Proteins extended) is an extended version of the SCOP database for classification of proteins based on structural relations. Astral compendium (21) provides sequences and coordinates for domains of 7 out of 12 classes of SCOPe. All sequences from Astral SCOPe version 2.08, filtered at 40% sequence identity, were used as query sequences to identify potential homologues. The NCBI Non-Redundant Database of protein sequences (NR, version Sep 2022) (22) was used to search for homologues. For domain identification in query sequences, the searches utilized the conserved domain database (23) (cdd_delta) from NCBI.

#### Sequence search

The DELTA-BLAST tool was employed for this purpose, using the BLAST executables version 2.13.0 (24). The searches utilised the conserved domain database (cdd_delta) from NCBI with a domain inclusion threshold of 0.05 for RPS-BLAST. An E-value threshold of 0.001 was set to consider a hit significant. The sequence database used for these searches was the NCBI NR (non-redundant) database from September 2022.

#### Validation

The hits identified from the DELTA-BLAST search, starting from individual PASS2 entries as queries, were combined for each superfamily. These sequence hits were validated using HMMs from PASS2 alignment and single-query HMMs built from Astral sequence set at 70% sequence identity. The HMMs (Hidden Markov models) from PASS2 version 2.4 and version 2.7 were used for validation. The sequences for building single query HMMs were obtained from scop.berkley.edu webserver. Single query HMMs were built using the hmmbuild program from the HMMER (version = 3.3.2) suite. HMMSCAN from the HMMER suite was used for validation with an E-value threshold of 0.01. The sequence hits from each superfamily were used as a query set. The HMMs from PASS2.4, PASS2.7 and single query HMMs from Astral sequence set at 70% sequence identity were compiled together and used as a target set. The domain architectures for the sequences were also obtained using the domain table output of the HMMSCAN results.

#### Identification of domain architectures

The domain architectures of identified homologues were determined using HMMSCAN, utilising the PASS2 HMM library along with single query HMMs. This library consists of PASS2 versions 4 (25) and 7 (26), which correspond to SCOPe versions 1.75 and 2.07, respectively and single query HMMs generated from members of SCOPe 2.08. The regions of each sequence that aligned with an HMM were annotated with the domain associated with the superfamily of that HMM. The Pfam domain architectures for the validated homologues were also identified using HMMSCAN and the Pfam-A HMM library.

#### Organisation of the website

The website for organising and analysing protein domain superfamilies across GenDiS database consists of two main components: the user end (front end) and the server end (backend). The front end was developed using HTML, CSS, JavaScript, bootstrap and jQuery, providing an intuitive and user-friendly interface and a dynamic and responsive website. AJAX was used for asynchronous requests to the server.

Data for the backend is stored using MongoDB, a flexible and scalable NoSQL database. The backend logic and webpage rendering are handled by Python Flask, a lightweight web framework. Data collection and storage involved pymongo for accessing the MongoDB database containing pre-collected data. Custom Python modules were used for parsing XML files and generating CSV files for visualisation purposes (27). The tool for domain architecture prediction utilises HMMSCAN from HMMER version 3.3.2, while the sequence alignment tools employ Clustal Omega version 1.2.4 (28).

#### Analysis using case studies

This study utilised a bioinformatics pipeline to investigate the taxonomic distribution and domain architectures of glycolysis-related enzymes. All KEGG (29) orthologies related to glycolysis were retrieved using the KEGG-API. This provided a comprehensive list of orthologous groups involved in glycolytic pathways. Using the obtained KEGG orthologies, all associated genes were identified through the KEGG- ORGANISMS database. This ensured the collection of genes that are directly linked to the orthologies involved in glycolysis across different organisms. The taxonomic distribution of the identified genes was analysed to understand the prevalence of these genes across various taxa. This involved categorising the genes according to their taxonomic affiliations using the KEGG-ORGANISMS database. The validated homologues for the identified genes were obtained from the GenDiS 3 database.

This step involved searching for homologous sequences using NCBI Accession IDs. The glycolysis enzymes were identified within the GenDiS database. The domain architecture of the identified glycolysis enzymes was fetched from the GenDiS 3 database using HMMSCAN results provided in the GenDiS database. The taxonomic distribution of the domain architectures was analysed to understand the evolutionary relationships and distribution patterns of the domain architectures across different taxa.

## Results

Using DELTA-BLAST to search for homologues, 15,128 members from 2,060 superfamilies were queried against the NCBI NR database (September 2022), which comprises 504,094,943 unique protein sequences. This search yielded 151,622,506 unique hits as potential homologues across all superfamilies (approximately 30% of the NR) (Table 1). Following validation with HMMSCAN, 116,393,303 of these sequences were confirmed as true positives (approximately 23% of the NR). The true positive rates across different superfamilies are distributed widely (please see Table ST2 and ST3 for list of superfamilies with highest and lowest true positive rates, respectively), with 50% of the superfamilies showing a true positive rate of more than 0.87 and 75% having rates of more than 0.65 for DELTA-BLAST hits (Fig 2).

**Fig 1:**
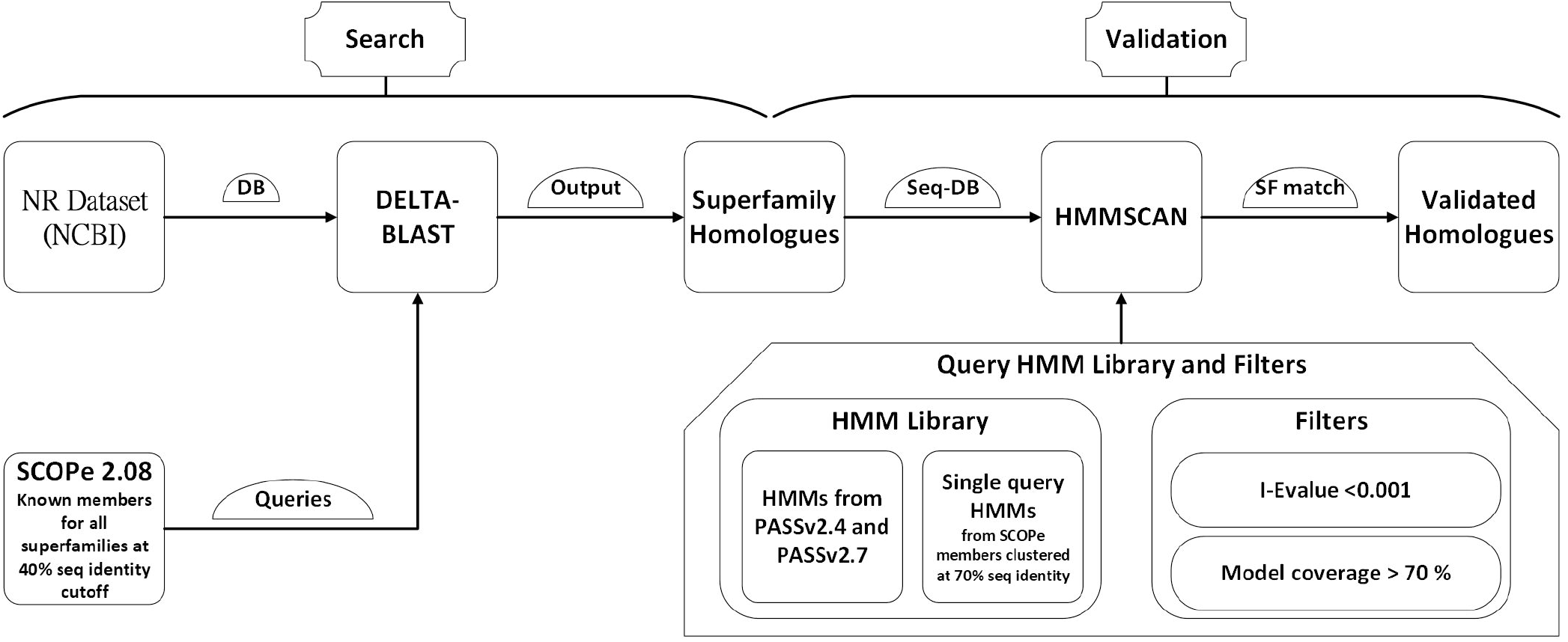
GenDiS3 workflow for search and validation of homologues.

**Table 1:** Total number of superfamilies, number of DELTABLAST hits, true positives after validation and mean true positive rate for each structural class of SCOPe.

| Class | Total #SFs | Total DELTABLAST hits | Total true positives | Mean True positive rate |
| --- | --- | --- | --- | --- |
| Mostly alpha (a) | 519 | 33,745,701 | 20,690,279 | 0.63694 |
| Mostly beta (b) | 374 | 36,009,271 | 21,973,707 | 0.60450 |
| Alpha and beta [a+b] (d) | 578 | 48,232,485 | 34,602,755 | 0.68532 |
| Alpha or beta [a/b] (c) | 247 | 67,224,602 | 53,394,941 | 0.77172 |
| Small proteins (g) | 73 | 5,315,902 | 3,726,663 | 0.66218 |
| Multi-domain proteins (e) | 130 | 5,867,215 | 3,991,323 | 0.67400 |
| Membrane proteins (f) | 139 | 6,003,912 | 4,053,609 | 0.67860 |

**Fig 2:**
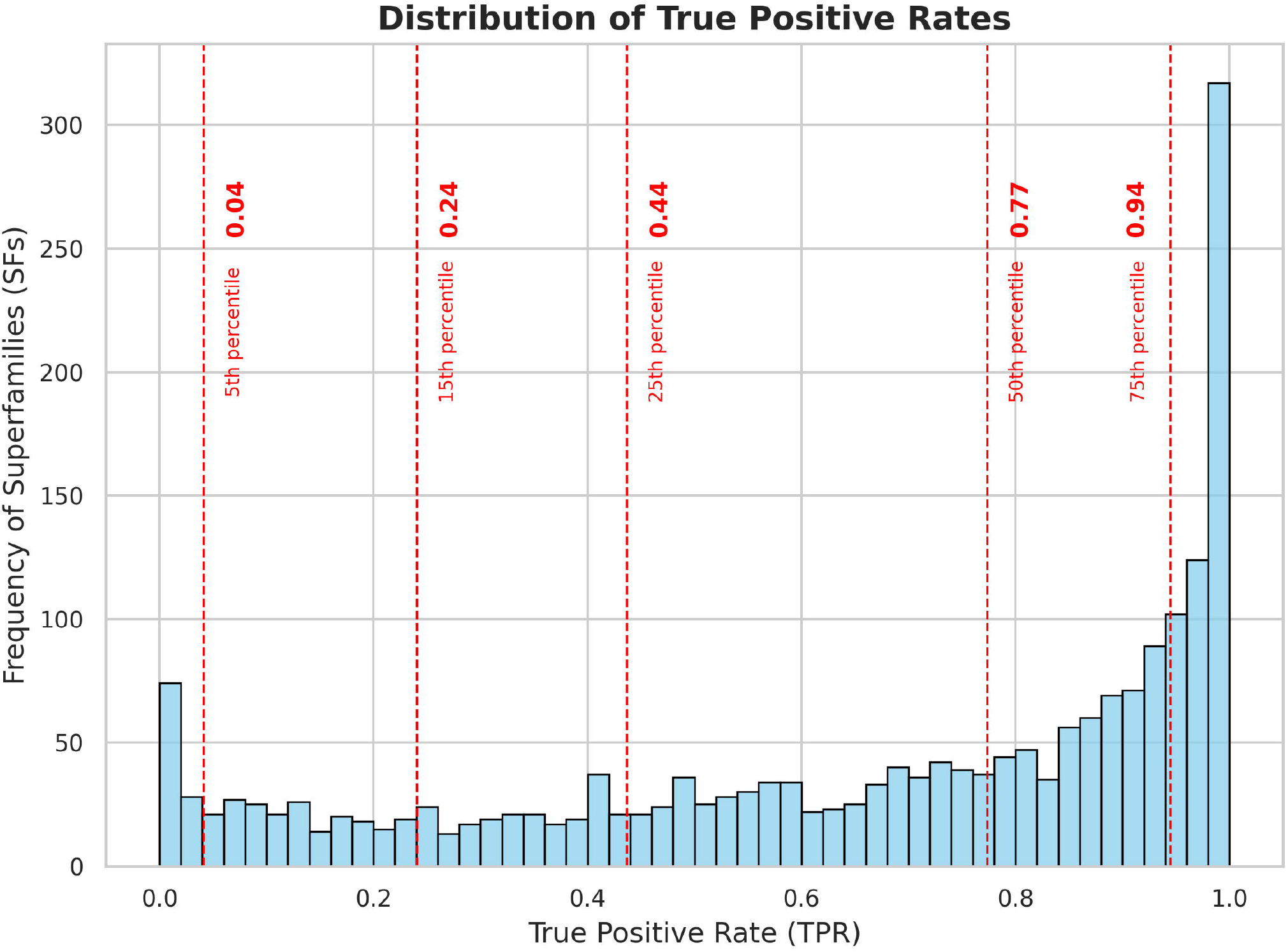
Frequency distribution of superfamilies with respect to true positive rates of the search strategy. The red lines indicate the 5th, 15th, 25th, 50th, and 75th percentiles of the data. From the plot we can observe that 50% of the superfamilies have a true positive rate of more than 0.77 for DELTA-BLAST, 75% of the superfamilies have a true positive rate of more than 0.44 for DELTA-BLAST.Analysis of domain architectures.

The delineation of domain architectures, in homologues where there is at least single presence of the central domain of interests, offers birds-eye view of the variety in biological processes where the central domain can get recruited. This also provides a comprehensive picture of the “social” or isolated nature of the central domain (30).

The superfamily c.37.1, identified as P-loop containing nucleoside triphosphate hydrolases (SF: 52540), is the most prevalent in the GenDiS database. This also corresponds to one of the superfolds (8). It is associated with the highest number of sequences, with the single domain architecture (52540) linked to approximately 2.5 million sequences. Additionally, the double domain architecture (52540∼52540) is found in 736 thousand sequences. The majority of homologues associated with this superfamily are from Eukaryotes, with a smaller proportion belonging to viruses.

The second most populated superfamily is c.2.1, known as NAD(P)-binding Rossmann-fold domains (SF: 51735). There are 1.69 million homologues associated with single domain architecture from this superfamily. Interestingly, around 500 thousand sequences from the 51735 superfamily also include the 6- phosphogluconate dehydrogenase C-terminal domain-like (SF: 48179), as a co- existing domain, with the combined architecture as 51735∼48179. This co-occurrence likely reflects a functional relationship, as both domains are involved in dehydrogenase activity where the Rossmann-fold domain binds cofactors like NAD(P)H, and the 6PGD domain catalyses specific metabolic reactions. All the other superfamilies are similarly associated predominantly with eukaryotic sequences.

The third highest populated superfamily is c.67.1, identified as PLP-dependent transferases (SF: 53383). The single domain architecture of this superfamily is associated with 1.6 million sequences.

Among the ten most prevalent domain architectures of homologues, nine consist of single domains, suggesting that single domain architectures are more common overall.

Indeed, the participation in multi-domain architectures is not uniform for all domains. Only a few domains have higher tendency to form multi- domain architectures. A graph theoretical approach of the study of domain architectures in GenDis3.0, where the nodes were individual domains and the edges represent co-existence of domains in a gene, revealed that the average degree of this graph was about 69. But some domains have a higher degree, implying that they are more likely to form multi-domain architectures. Domains from Superfamily c.37.1, P-loop containing nucleoside triphosphate hydrolases, have the highest degree of 1244 along with being the most populated superfamily.

Table 2 lists the SCOPe codes and corresponding superfamily names of protein domains that show the highest degree of coexistence with other domains in multi-domain architectures. The degree of coexistence is indicated by the number of unique domain combinations in which each domain is found. The data highlights domains such as P-loop containing nucleoside triphosphate hydrolases (c.37.1), NAD(P)-binding Rossmann-fold domains (c.2.1), and Protein kinase-like (PK-like) (d.144.1) as the most frequently co-occurring, suggesting their widespread involvement in diverse biological functions.

**Table 2:** Domains with the Highest Degree of Coexistence in multi-domain Architectures.

| **SCOPe Code** | **Superfamily name** | **Degree** |
| --- | --- | --- |
| c.37.1 | P-loop containing nucleoside triphosphate hydrolases | 1244 |
| c.2.1 | NAD(P)-binding Rossmann-fold domains | 919 |
| d.144.1 | Protein kinase-like (PK-like) | 911 |
| c.69.1 | alpha/beta-Hydrolases | 735 |
| a.4.5 | Winged helix' DNA-binding domain | 728 |
| c.47.1 | Thioredoxin-like | 714 |
| b.40.4 | Nucleic acid-binding proteins | 670 |
| d.58.7 | RNA-binding domain, RBD, aka RNA recognition motif (RRM) | 652 |
| b.69.4 | WD40 repeat-like | 641 |
| d.211.1 | Ankyrin repeat | 629 |
| c.66.1 | S-adenosyl-L-methionine-dependent methyltransferases | 620 |
| c.108.1 | HAD-like | 598 |
| a.118.8 | TPR-like | 596 |
| g.44.1 | RING/U-box | 592 |
| a.4.1 | Homeodomain-like | 581 |
| c.1.8 | (Trans)glycosidases | 574 |
| c.67.1 | PLP-dependent transferases | 559 |
| g.37.1 | beta-beta-alpha zinc fingers | 549 |
| a.39.1 | EF-hand | 546 |
| c.55.1 | Actin-like ATPase domain | 529 |
| c.55.3 | Ribonuclease H-like | 510 |
| c.3.1 | FAD/NAD(P)-binding domain | 497 |
| d.3.1 | Cysteine proteinases | 493 |
| b.1.1 | Immunoglobulin | 490 |
| c.10.2 | L domain-like | 489 |

### Database Browsing and Visualisation

The updated GenDiS3 database offers hierarchical browsing of superfamily hits through the SCOPe classification, including folds and classes. This feature facilitates user navigation and discovery. For each superfamily, users can view SCOPe and Pfam domain architectures of the homologues, along with the frequency of each domain architecture within the superfamily. This functionality provides valuable insights into the structural diversity within superfamilies.

### Taxonomic Distribution and Visualisation

An interactive sunburst chart visualises the taxonomic distribution of superfamily hits, offering an intuitive way to explore the evolutionary spread of the identified homologues This visualisation aids in understanding the taxonomic breadth and evolutionary context of the superfamilies (Fig 3a).

**Fig 3:**
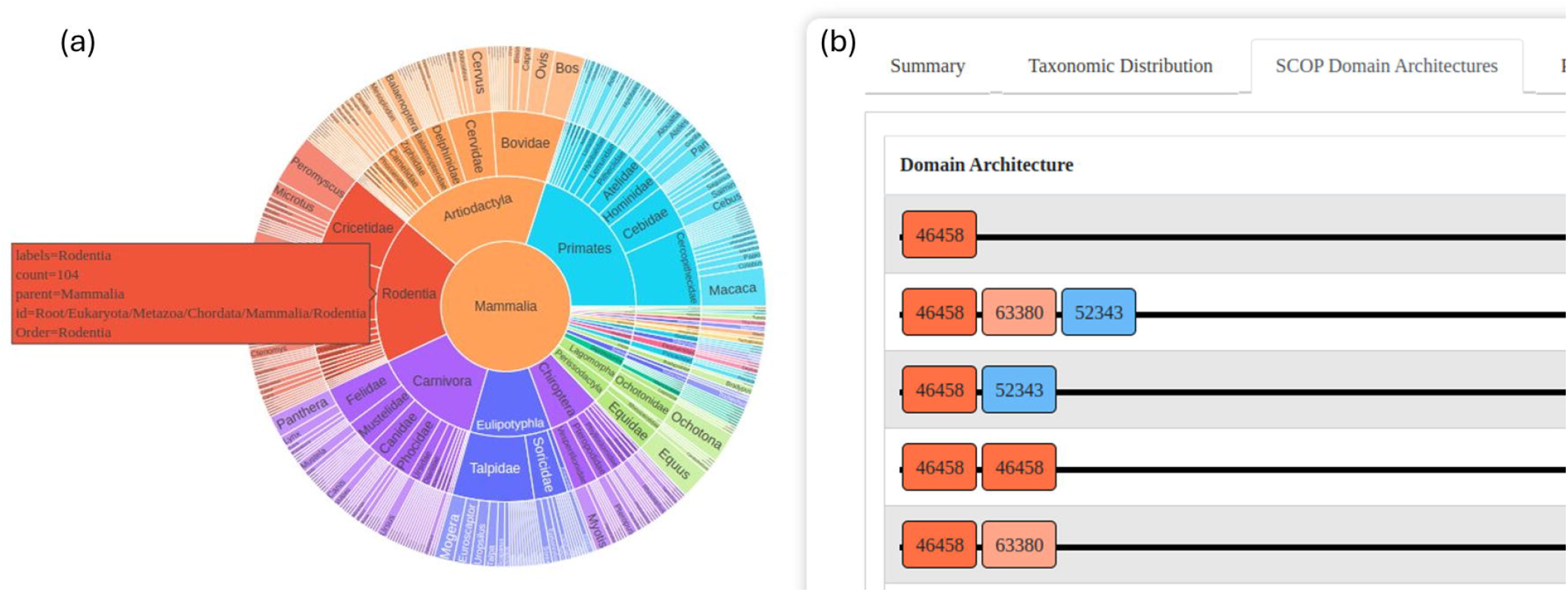
The GenDiS3 web interface for visualising taxonomic distribution and domain architectures. (a) Sunburst chart (interactive) showing the taxonomic distribution of a superfamily. (b) SCOP domain architectures of a superfamily.

### Tools and capabilities

#### Domain architecture prediction tools

This tool allows users to predict the domain architecture of their provided sequences. Users can choose to obtain either SCOPe domain architecture (DA) or Pfam domain architecture (DA). This capability is crucial for researchers looking to annotate new sequences or verify the domain structures of known proteins, facilitating functional annotation and evolutionary studies.

#### Align two genomes of a given superfamily

This tool aligns superfamily hits of a specific superfamily from two different genomes. It helps in comparative genomics studies by allowing researchers to examine the evolutionary conservation and divergence of superfamilies between two species. This tool is particularly useful for identifying conserved domains and understanding the functional implications of superfamily distribution across different organisms.

#### Align Input Sequence with Genome of a Given Superfamily

Similar to the previous tool, this one aligns a user-provided sequence with the genome of a given superfamily. It enables researchers to investigate how a specific sequence aligns with known superfamily members within a genome, aiding in the identification of homologous regions and potential functional elements within the genome. This tool is beneficial for pinpointing evolutionary relationships and functional annotations of user-specific sequences.

#### Blast search superfamily homologues

This tool allows users to search for similar sequences within all the homologues of a superfamily using a query sequence. The tool uses blastp executable from NCBI. The tool will help researchers to look for genes within superfamilies using reference sequences.

### Case Study 1: The glycolysis pathway in GenDiS

Glycolysis is a fundamental metabolic pathway found in nearly all living organisms, from bacteria to humans. It involves a diverse set of enzymes with varying structures and functions. In this study, we explored the taxonomic distribution of various SCOPe domains within glycolysis pathway enzymes using the GenDiS database. To identify these enzymes, we mapped known glycolysis enzymes to their respective KEGG orthology (KO) identifiers via KEGG. These KO identifiers were then used to query GenDiS for homologues containing the relevant SCOPe domains (Fig. S1). Here are some notable observations:

### Seeing the knowns: phosphofructokinase

The KEGG pathway for glycolysis includes two orthologies for phosphofructokinase: 6-phosphofructokinase (K24182) and ADP-dependent phosphofructokinase (K00918). The 6-phosphofructokinase (6-PFK) is primarily observed in bacteria, while ADP-dependent phosphofructokinase is predominantly present in archaea (Fig. S2). This observation aligns with expected patterns, as bacteria typically utilise ATP-dependent 6-phosphofructokinase in classical glycolysis pathways (31). In contrast, archaea, often adapted to extreme environments, employ ADP-dependent phosphofructokinase to maintain metabolic flexibility (32).

### Hexokinase

Hexokinase is observed to be present in all kingdoms, emphasising its essential role in glycolysis. The most frequently found SCOPe domain architectures in these proteins are two domain and four domain repeats of 53067 (Actin-like ATPase domain superfamily containing the hexokinase family). All kingdoms, except animals, typically exhibit a two tandem repeat domain architecture, whereas animals display a four tandem repeat domain architecture (Fig. S3).

### Enolase

Enolase is present across all kingdoms, demonstrating its fundamental role in glycolysis. The most common domain architecture identified was 54826∼51604 (Enolase NTD∼CTD) (Fig. S4). Additionally, it was notable that some enolase proteins also coexisted with the HLH DNA-binding domain (47459), pointing to alpha-enolases which are nuclear and found to regulate gene expression (33). This also exemplifies the analysis of domain architectures could reveal potential multi- functional role of central domains.

The analysis of glycolysis enzymes across different kingdoms revealed distinct patterns in their taxonomic distribution and domain architectures (Fig 4). Phosphofructokinase showed a clear divergence, with 6-phosphofructokinase predominantly found in bacteria and ADP-dependent phosphofructokinase mainly in archaea, reflecting their metabolic adaptations. Enolase was universally present across all kingdoms, with the most common domain architecture being Enolase NTD∼CTD, and some proteins also exhibiting the HLH DNA-binding domain, suggesting additional functional roles. Hexokinase, also present in all kingdoms, displayed varying domain architectures: two tandem repeats in most kingdoms and four tandem repeats in animals, highlighting a possible evolutionary expansion in animals. These findings show the evolutionary diversity and specialisation of glycolytic enzymes, driven by the metabolic needs and environmental adaptations of different organisms.

**Fig 4:**
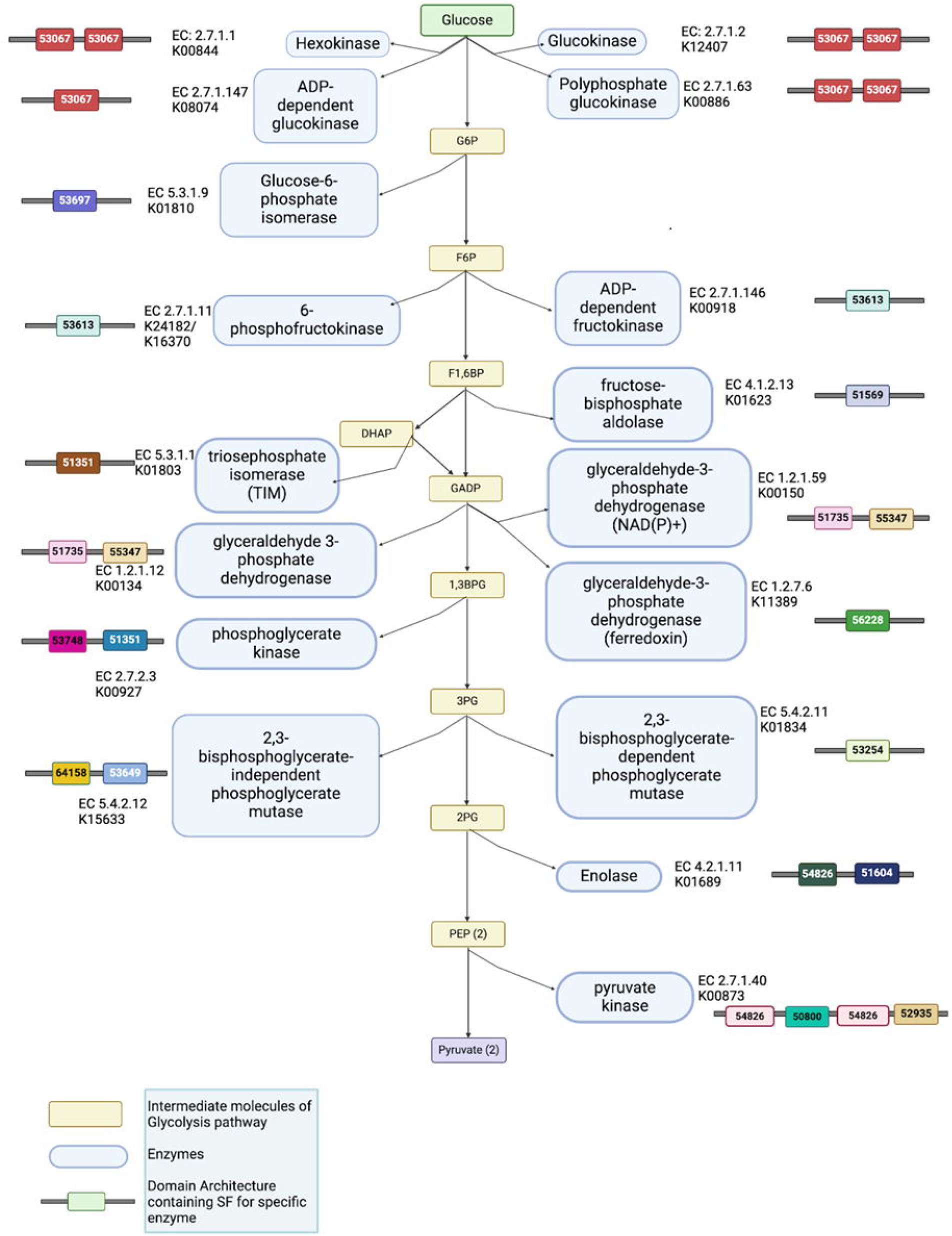
The glycolysis pathway. Enzymes are represented by light blue boxes; yellow boxes correspond to intermediate molecules within the pathway. Colourful boxes flanking the pathway represent the domain architecture of each enzyme where each SCOP domain is represented by the superfamily code. SF 53067: c.55.1 (Actin-like ATPase domain), SF 53697: c.80.1 (SIS domain), SF 53613: c.72.1 (Ribokinase-like), SF 51569: c.1.10 (Aldolase), SF 51351: c.1.1 (Triosephosphate isomerase (TIM)), SF 51735: c.2.1 (NAD(P)-binding Rossmann-fold domains), SF 55347: d.81.1 (Glyceraldehyde-3-phosphate dehydrogenase-like, C-terminal domain), SF 53748: c.86.1 (Phosphoglycerate kinase), SF 56228: d.152.1 (Aldehyde ferredoxin oxidoreductase, N-terminal domain), SF 64158: c.105.1 (2,3-Bisphosphoglycerate-independent phosphoglycerate mutase, substrate-binding domain), SF 53649: c.76.1 (Alkaline phosphatase-like), SF 53254: c.60.1 (Phosphoglycerate mutase-like), SF 54826: d.54.1 (Enolase N-terminal domain-like), SF 51604: c.1.11 (Enolase C-terminal domain-like), SF 50800: b.58.1 (PK beta-barrel domain-like), SF 52935: c.49.1 (PK C-terminal domain-like).

### Case Study 2: LOG Gene implicated in the formation of prickles

Prickles, small, sharp outgrowths on the stems and leaves of many plants, serve as a natural defence mechanism against herbivores. While various morphological features and underlying genetic factors contributing to prickle formation have been identified, recent studies have highlighted the role of the LOG (Lonely Guy) gene as a key player in this process (34). The LOG gene codes for cytokinin riboside 5’- monophosphate phosphoribohydrolase (CRMP) enzyme. It was initially known for its role in cytokinin biosynthesis and has been implicated in the formation of prickles across a wide range of plant species, including economically important crops such as eggplant and rose. James W. Satterlee et al. showed that mutations in the LOG gene can result in a prickle-less phenotype in *Solanum* sp. Some of these mutations are mapped onto the structure of CRMP from *Arabidopsis thaliana* (Fig 5). Interestingly, even the intronic mutations are mapped on the peripheral helices, away from the central beta-sheet and the active site region.

**Fig 5:**
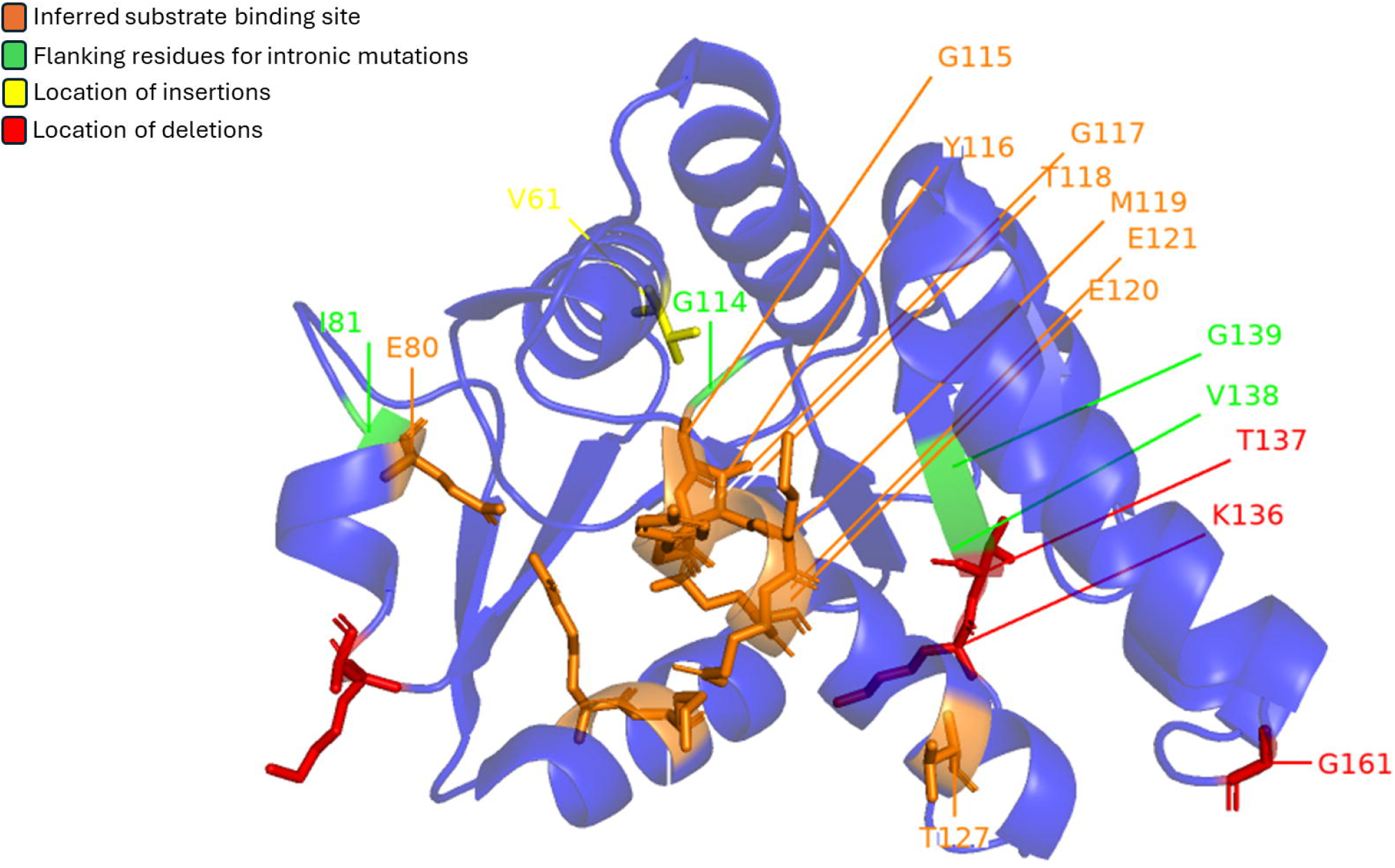
Mutations found in LOG genes from *Solanum* genus mapped on Cytokinin riboside 5’-monophosphate phosphoribohydrolase (LOG8) from *Arabidopsis thaliana* (PDB: 1YDH). Red represents location of deletions, yellow represents location of insertions and green marks the flanking residues for intronic mutations. The orange area marks the inferred substrate binding site.

We know that CRMP (encoded by the LOG gene) belongs to the MCP/YpsA-like superfamily as classified by SCOPe. We searched for all the homologues of LOG gene in MCP/YpsA-like superfamily using the built-in blastp search tool of GenDis3.0 database, with LOG genes from *A. thaliana* and *O. sativa* (from UniProt) as queries. From this search, we found 5,812 homologues for the LOG gene protein belonging to Viridiplantae kingdom. Over a thousand of these are unannotated hypothetical proteins and this pipeline can be used to annotate these as LOG genes.

### Enhancing Homologue Detection Accuracy

The use of DELTA-BLAST and HMMSCAN in identifying homologues has significantly improved the accuracy and reliability of the GenDiS database. DELTA-BLAST’s integration of domain-specific information allows for more sensitive searches, while HMMSCAN provides robust validation by comparing identified sequences against well-curated profile HMM databases. The results show that this approach yields a higher true positive rate across various structural classes, particularly for superfamilies with conserved domain architectures. These findings suggest that the updated GenDiS database is a powerful resource for the classification and annotation of protein domains, which is crucial for both functional and evolutionary studies. One of the primary challenges addressed in this study is the exponential growth of protein sequence data, which far outpaces the determination of their corresponding structures. The updated GenDiS database helps to mitigate this issue by providing a more efficient framework for linking sequences to known structures. By enhancing the accuracy of domain architecture predictions and facilitating the identification of distant homologues, the GenDiS database plays a critical role in expanding our understanding of protein structures and their functional implications, even in the absence of experimental structure determination.

### Understanding Evolutionary Relationships and Structural Diversity

The taxonomic analysis of glycolysis-related enzymes, including phosphofructokinase, revealed patterns consistent with established knowledge. For example, ATP-dependent phosphofructokinase is well-documented in bacteria, while ADP-dependent phosphofructokinase (ADP-PFK) is more commonly found in archaea. These findings align with previous research and reflect the metabolic adaptations of these organisms to their environments. Similarly, the variation in hexokinase domain architectures across kingdoms shows the evolutionary pressures that have shaped these enzymes to fulfil essential metabolic roles. These insights provide a deeper understanding of how protein domain architectures evolve in response to functional demands and environmental conditions.

### Complexity of Multi-Domain Architectures

The study also delved into the complexity of multi-domain architectures, which are prevalent in many proteins and critical for their multi-functionality. The analysis revealed that whereas single domain architectures are common, certain domains, particularly those from the c.37.1 superfamily, have a high propensity to form multi-domain architectures. This finding highlights the importance of such “social” domains (30) in facilitating diverse biological functions. Understanding the tendencies of specific domains to participate in multi-domain architectures can provide valuable insights into their evolutionary history and functional roles.

## Conclusion

The study demonstrates the importance of advanced computational methodologies in enhancing protein domain classification and mutations which provide valuable insights into the evolutionary trends of important proteins. While the updated GenDiS database represents a significant advancement in protein domain classification and evolutionary studies, several challenges remain. Despite the rapid and robust nature of DELTA-BLAST, the mere runs on such sequence searches require months (please see GenDis3.0 about-timeframe webpage) (Table ST1). The scalability of the database will acquire prime importance since protein sequence data continues to grow and how the computational resources respond to this ever-growing primary data will be crucial for maintaining its utility. Future updates will need to focus on optimising data management, avoidance of validation step and processing capabilities to handle this influx efficiently.

This study has demonstrated the effectiveness of advanced bioinformatics tools in enhancing the classification and annotation of protein domains within the GenDiS database. By addressing key challenges in homologue detection, sequence data growth, and domain architecture complexity, the updated database offers a valuable resource for understanding the structural and functional diversity of proteins.

Continued development and expansion of the GenDiS database will be essential for keeping pace with the rapid advancements in genomics and structural biology, ultimately contributing to a deeper understanding of protein function and evolution. It also offers a powerful resource for researchers in functional annotation and evolution of protein domain superfamilies.

## Acknowledgements

RS acknowledges funding and support provided by JC Bose Fellowship (JBR/2021/000006) from Science and Engineering Research Board, India and Bioinformatics Centre Grant funded by Department of Biotechnology, India (BT/PR40187/BTIS/137/9/2021). RS would also like to thank Institute of Bioinformatics and Applied Biotechnology for the funding through her Mazumdar-Shaw Chair in Computational Biology (IBAB/MSCB/182/2022).

## Supplementary Data

**Table ST1:** Time required for the search and validation runs.

**Table ST2:** Superfamilies with the highest true positive rate (top 50).

**Table ST3:** Superfamilies with lowest true positive rates (bottom 50).

**Fig S1:** Workflow for identification of Glycolysis enzymes in GenDiS using KEGG Pathways database.

**Fig S2:** Plot showing number of homologues for ADP-dependent fructokinase and 6- phosphofructokinase and their abundance in archaea and bacteria. Crystal structures of 6-phosphofructokinase (A 2abq:A) and ADP-dependent fructokinase (B 1u2x:A).

**Fig S3:** Kingdom wise distribution of enolase proteins and frequency of various domain architectures found in hexokinase.

**Fig S4**: Kingdom wise distribution of enolase proteins and frequency of various domain architectures found in enolase.

