## Supplementary Figures and Tables for "GenDiS3 database: census on prevalence of protein domain superfamilies of known structure in the entire sequence database"

### Supplementary Data

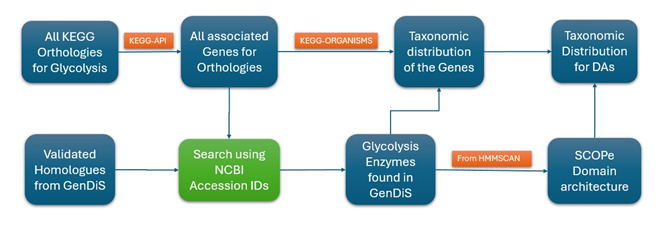

**Fig S1:** Workflow for identification of Glycolysis enzymes in GenDiS using KEGG Pathways database.

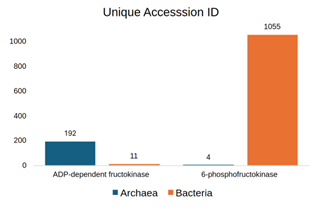

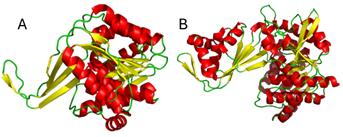

**Fig S2:** Plot showing number of homologues for ADP-dependent fructokinase and 6-phosphofructokinase and their abundance in archaea and bacteria. Crystal structures of 6-phosphofructokinase (A 2abq:A) and ADP-dependent fructokinase (B 1u2x:A).

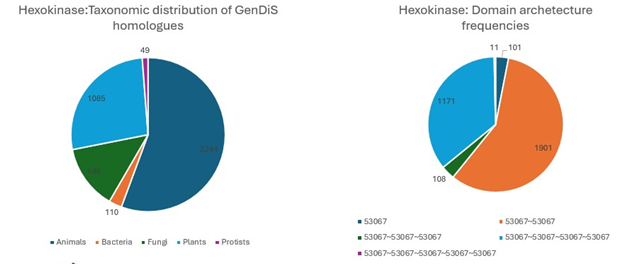

**Fig S3:** Kingdom wise distribution of enolase proteins and frequency of various domain architectures found in hexokinase.

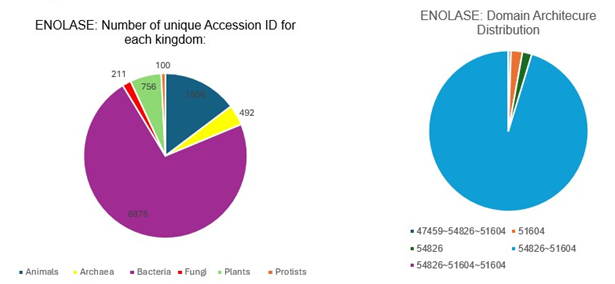

**Fig S4**: Kingdom wise distribution of enolase proteins and frequency of various domain architectures found in enolase.

**Table ST1:** Time required for the search and validation runs.

| **Class** | **Total #SFs** | **Total DELTA-BLAST hits** | **Total Validated Hits** | **Time Required for DELTA-BLAST** | **Time Required for HMMSCAN** |
| --- | --- | --- | --- | --- | --- |
| Mostly alpha | 519 | 33M | 20M | ~4000H | ~624H |
| Mostly beta | 374 | 36M | 21M | ~3500H | ~984H |
| Alpha and beta | 578 | 48M | 34M | ~1900H | ~840H |
| Alpha or beta | 247 | 67M | 53M | ~3600H | ~1512H |
| Small proteins | 73 | 5M | 3M | ~500H | ~260H |
| Multidomain proteins | 130 | 5M | 3M | ~620H | ~380H |
| Membrane proteins | 139 | 6M | 4M | ~500H | ~400H |
| **Total** | **2060** | **202M** | **142M** | **~6 months** | **~4 months** |

**Table ST2:** Superfamilies with the highest true positive rate (top 50).

| **sf** | **sf_code** | **class** | **description** | **tpr** |
| --- | --- | --- | --- | --- |
| 142695 | c.144.1 | Alpha and beta (a/b) | RibA-like | 1 |
| 57938 | g.54.1 | Small proteins | DnaJ/Hsp40 cysteine-rich domain | 1 |
| 51713 | c.1.20 | Alpha and beta (a/b) | tRNA-guanine transglycosylase | 1 |
| 55271 | d.75.2 | Alpha and beta (a+b) | DNA repair protein MutS, domain I | 1 |
| 52255 | c.23.8 | Alpha and beta (a/b) | N5-CAIR mutase (phosphoribosylaminoimidazole carboxylase, PurE) | 1 |
| 54821 | d.53.1 | Alpha and beta (a+b) | Ribosomal protein S3 C-terminal domain | 1 |
| 52021 | c.8.3 | Alpha and beta (a/b) | Carbamoyl phosphate synthetase, small subunit N-terminal domain | 1 |
| 82051 | b.117.1 | All beta | Obg GTP-binding protein N-terminal domain | 1 |
| 81886 | a.172.1 | All alpha | Helical scaffold and wing domains of SecA | 1 |
| 141259 | b.34.18 | All beta | CarD-like | 1 |
| 54364 | d.15.8 | Alpha and beta (a+b) | Translation initiation factor IF3, N-terminal domain | 1 |
| 75620 | e.38.1 | Multi-domain proteins | Release factor | 1 |
| 159936 | d.15.14 | Alpha and beta (a+b) | NSP3A-like | 1 |
| 54810 | d.52.2 | Alpha and beta (a+b) | GMP synthetase C-terminal dimerisation domain | 1 |
| 100920 | b.130.1 | All beta | Heat shock protein 70kD (HSP70), peptide-binding domain | 1 |
| 55973 | d.130.1 | Alpha and beta (a+b) | S-adenosylmethionine synthetase | 1 |
| 47644 | a.46.1 | All alpha | Methionine synthase domain | 1 |
| 53178 | c.56.3 | Alpha and beta (a/b) | Peptidyl-tRNA hydrolase-like | 1 |
| 47446 | a.36.1 | All alpha | Signal peptide-binding domain | 1 |
| 56712 | e.10.1 | Multi-domain proteins | Prokaryotic type I DNA topoisomerase | 0.999988 |
| 47741 | a.56.1 | All alpha | CO dehydrogenase ISP C-domain like | 0.999986 |
| 82714 | d.225.1 | Alpha and beta (a+b) | Multidrug efflux transporter AcrB TolC docking domain; DN and DC subdomains | 0.999985 |
| 143076 | d.302.1 | Alpha and beta (a+b) | Coronavirus NSP8-like | 0.999985 |
| 144246 | g.86.1 | Small proteins | Coronavirus NSP10-like | 0.999985 |
| 140367 | a.8.9 | All alpha | Coronavirus NSP7-like | 0.999984 |
| 55200 | d.68.1 | Alpha and beta (a+b) | Translation initiation factor IF3, C-terminal domain | 0.999983 |
| 103642 | g.74.1 | Small proteins | Sec-C motif | 0.999982 |
| 101904 | b.68.10 | All beta | GyrA/ParC C-terminal domain-like | 0.999982 |
| 55186 | d.67.1 | Alpha and beta (a+b) | ThrRS/AlaRS common domain | 0.999982 |
| 101816 | b.140.1 | All beta | Replicase NSP9 | 0.999982 |
| 64005 | c.101.1 | Alpha and beta (a/b) | Undecaprenyl diphosphate synthase | 0.999982 |
| 53671 | c.78.1 | Alpha and beta (a/b) | Aspartate/ornithine carbamoyltransferase | 0.999981 |
| 140490 | a.29.12 | All alpha | Nqo1C-terminal domain-like | 0.999981 |
| 74982 | b.111.1 | All beta | Small protein B (SmpB) | 0.99998 |
| 55040 | d.58.21 | Alpha and beta (a+b) | Molybdenum cofactor biosynthesis protein C, MoaC | 0.99998 |
| 140990 | a.269.1 | All alpha | FtsH protease domain-like | 0.99998 |
| 46767 | a.4.2 | All alpha | Methylated DNA-protein cysteine methyltransferase, C-terminal domain | 0.99998 |
| 48024 | a.81.1 | All alpha | N-terminal domain of DnaB helicase | 0.999979 |
| 52738 | c.40.1 | Alpha and beta (a/b) | Methylesterase CheB, C-terminal domain | 0.999976 |
| 160099 | d.346.1 | Alpha and beta (a+b) | SARS Nsp1-like | 0.999975 |
| 56420 | d.167.1 | Alpha and beta (a+b) | Peptide deformylase | 0.999973 |
| 81345 | f.22.1 | Membrane and cell surface proteins | ABC transporter involved in vitamin B12 uptake, BtuC | 0.999972 |
| 54368 | d.15.9 | Alpha and beta (a+b) | Glutamine synthetase, N-terminal domain | 0.99997 |
| 63882 | b.103.1 | All beta | MoeA N-terminal region -like | 0.999968 |
| 69864 | d.210.1 | Alpha and beta (a+b) | Argininosuccinate synthetase, C-terminal domain | 0.999967 |
| 48334 | a.113.1 | All alpha | DNA repair protein MutS, domain III | 0.999966 |
| 56047 | d.140.1 | Alpha and beta (a+b) | Ribosomal protein S8 | 0.999966 |
| 55594 | d.94.1 | Alpha and beta (a+b) | HPr-like | 0.999964 |
| 69765 | d.79.5 | Alpha and beta (a+b) | IpsF-like | 0.99996 |
| 75625 | e.39.1 | Multi-domain proteins | YebC-like | 0.999959 |

**Table ST3:** Superfamilies with lowest true positive rates (bottom 50).

| **sf** | **sf_code** | **class** | **description** | **tpr** |
| --- | --- | --- | --- | --- |
| 75412 | d.214.1 | Alpha and beta (a+b) | Hypothetical protein MTH1880 | 0.004928 |
| 160887 | d.377.1 | Alpha and beta (a+b) | Rv2827c C-terminal domain-like | 0.004659 |
| 158842 | a.296.1 | All alpha | PMT central region-like | 0.004633 |
| 49498 | b.5.1 | All beta | alpha-Amylase inhibitor tendamistat | 0.004199 |
| 50012 | b.31.1 | All beta | EV matrix protein | 0.004136 |
| 69070 | a.150.1 | All alpha | Anti-sigma factor AsiA | 0.00311 |
| 140496 | a.30.6 | All alpha | HP1531-like | 0.003086 |
| 81986 | b.1.20 | All beta | Tp47 lipoprotein, middle and C-terminal domains | 0.002903 |
| 50934 | b.67.1 | All beta | Tachylectin-2 | 0.002436 |
| 64043 | c.102.1 | Alpha and beta (a/b) | Cell-division inhibitor MinC, N-terminal domain | 0.002354 |
| 103383 | e.45.1 | Multi-domain proteins | Antivirulence factor | 0.002269 |
| 111057 | d.274.1 | Alpha and beta (a+b) | Hypothetical protein PF0899 | 0.0021 |
| 140860 | a.118.24 | All alpha | Pseudo ankyrin repeat-like | 0.001932 |
| 111265 | d.281.1 | Alpha and beta (a+b) | Hemolytic lectin CEL-III, C-terminal domain | 0.001877 |
| 51156 | b.80.2 | All beta | Insect cysteine-rich antifreeze protein | 0.001825 |
| 116942 | a.232.1 | All alpha | RNA-binding protein She2p | 0.001399 |
| 159612 | c.52.4 | Alpha and beta (a/b) | TBP-interacting protein-like | 0.001177 |
| 101424 | a.118.20 | All alpha | Hypothetical protein ST1625 | 0.000932 |
| 254122 | f.60.1 | Membrane and cell surface proteins | Anthrax protective antigen C-terminal-like | 0.00066 |
| 144129 | g.84.1 | Small proteins | Vanabin-like | 0.000522 |
| 116965 | a.234.1 | All alpha | Hypothetical protein MPN330 | 0.000512 |
| 160761 | d.374.1 | Alpha and beta (a+b) | TTHC002-like | 0.000382 |
| 140979 | a.267.1 | All alpha | Topoisomerase V catalytic domain-like | 0.000143 |
| 101173 | a.38.2 | All alpha | Docking domain B of the erythromycin polyketide synthase (DEBS) | 0.000122 |
| 89428 | b.126.1 | All beta | Adsorption protein p2 | 9.91E-05 |
| 103589 | g.71.1 | Small proteins | Mini-collagen I, C-terminal domain | 0 |
| 418753 | d.58.64 | Alpha and beta (a+b) | Cytosolic domain of anoctamin channel-like | 0 |
| 418762 | d.399.1 | Alpha and beta (a+b) | Ebola nucleoprotein C-terminal domain-like | 0 |
| 418755 | d.58.65 | Alpha and beta (a+b) | DUF4478 domain-like | 0 |
| 418741 | d.397.1 | Alpha and beta (a+b) | Influenza virus PB2 cap binding domain-like | 0 |
| 418769 | d.400.1 | Alpha and beta (a+b) | Lem10 N-terminal domain-like | 0 |
| 418767 | d.230.9 | Alpha and beta (a+b) | P2X receptor transmembrane region | 0 |
| 418765 | d.230.8 | Alpha and beta (a+b) | P2X receptor head domain-like | 0 |
| 418740 | d.396.1 | Alpha and beta (a+b) | Influenza virus PB2 '627' domain-like | 0 |
| 418752 | d.58.63 | Alpha and beta (a+b) | Cytosolic domain of OSCA channel-like | 0 |
| 48686 | a.137.7 | All alpha | Proteinase A inhibitor IA3 | 0 |
| 418758 | d.398.1 | Alpha and beta (a+b) | NUC domain from CRISPR-associated protein Cas12a / Cpf1-like | 0 |
| 418770 | d.401.1 | Alpha and beta (a+b) | Cas3 C-terminal domain-like | 0 |
| 418772 | d.402.1 | Alpha and beta (a+b) | B domain from Cmr1-like | 0 |
| 418764 | d.230.7 | Alpha and beta (a+b) | Oligosaccharide binding protein | 0 |
| 418774 | d.403.1 | Alpha and beta (a+b) | Lewis(b) antigen binding domain of BabA-like | 0 |
| 103436 | f.23.24 | Membrane and cell surface proteins | PetL subunit of the cytochrome b6f complex | 0 |
| 161148 | g.2.4 | Small proteins | VhTI-like | 0 |
| 161029 | f.23.34 | Membrane and cell surface proteins | Photosystem II reaction center protein T, PsbT | 0 |
| 267600 | f.23.41 | Membrane and cell surface proteins | Photosystem II reaction center protein ycf12 | 0 |
| 418737 | g.102.1 | Small proteins | Jumonji C-terminal domain-like | 0 |
| 103451 | f.23.27 | Membrane and cell surface proteins | PetN subunit of the cytochrome b6f complex | 0 |
| 81552 | f.23.20 | Membrane and cell surface proteins | Subunit PsaX of photosystem I reaction centre | 0 |
| 418745 | d.15.15 | Alpha and beta (a+b) | Phosphoinositide 3-kinase alpha adaptor-binding domain (ABD)-like | 0 |
| 90148 | g.3.18 | Small proteins | DPY module | 0 |
